# Visual and auditory cortices represent acoustic speech-related information during silent lip reading

**DOI:** 10.1101/2022.02.21.481292

**Authors:** Felix Bröhl, Anne Keitel, Christoph Kayser

**Affiliations:** Department for Cognitive Neuroscience, Faculty of Biology, Bielefeld University, Universitätsstr. 25, 33615, Bielefeld, Germany; Psychology, University of Dundee, Scrymgeour Building, Dundee DD1 4HN, UK

**Keywords:** Speech entrainment, lip-reading, audio-visual, speech tracking, language, MEG

## Abstract

Speech is an intrinsically multisensory signal and seeing the speaker’s lips forms a cornerstone of communication in acoustically impoverished environments. Still, it remains unclear how the brain exploits visual speech for comprehension and previous work debated whether lip signals are mainly processed along the auditory pathways or whether the visual system directly implements speech-related processes. To probe this question, we systematically characterized dynamic representations of multiple acoustic and visual speech-derived features in source localized MEG recordings that were obtained while participants listened to speech or viewed silent speech. Using a mutual-information framework we provide a comprehensive assessment of how well temporal and occipital cortices reflect the physically presented signals and speech-related features that were physically absent but may still be critical for comprehension. Our results demonstrate that both cortices are capable of a functionally specific form of multisensory restoration: during lip reading both reflect unheard acoustic features, with occipital regions emphasizing spectral information and temporal regions emphasizing the speech envelope. Importantly, the degree of envelope restoration was predictive of lip reading performance. These findings suggest that when seeing the speaker’s lips the brain engages both visual and auditory pathways to support comprehension by exploiting multisensory correspondences between lip movements and spectro-temporal acoustic cues.

**Highlights:**

- Visual and auditory cortex represent unheard acoustic information during lip reading
- Auditory cortex emphasizes the acoustic envelope
- Visual cortex emphasizes a pitch signature
- Tracking of unheard features in auditory cortex is associated with behavior

## 1. Introduction

Speech is an intrinsically multisensory stimulus that is often conveyed via both acoustic and visual signals. Visual speech contains information that becomes particularly important in circumstances when the acoustic signal is impoverished, such as by background noises or distractors (Ross et al., 2007; Sumby and Pollack, 1954). In these cases listeners typically look at the speaker’s face to achieve a genuine multisensory benefit for comprehension, which in the brain is mediated by an enhanced cortical encoding of acoustic and phonemic speech features (Giordano et al., 2017; Mégevand et al., 2020; O’Sullivan et al., 2017; Zion Golumbic et al., 2013). However, in situations where only visual speech cues are available, i.e. during silent lip reading, visual signals allow comprehension also when the respective acoustic information is absent (Besle et al., 2008; Calvert et al., 1997; Calvert and Campbell, 2003; Grant and Seitz, 2000).

How exactly the brain represents the information derived from visual speech and how it exploits this for comprehension remains debated. One possibility is that visual speech is represented in regions of the auditory pathways, possibly exploiting speech-specific processes of the auditory system. Neuroimaging studies support this view by demonstrating the activation of the auditory cortex when participants view the articulation of words or pseudo-words, but not when viewing non-speech gestures (Bernstein et al., 2002; Besle et al., 2008; Calvert et al., 1997; Calvert and Campbell, 2003; Pekkola et al., 2005; Sams et al., 1991). Along this line, a recent study has suggested that the auditory cortex can reflect the unheard acoustic envelope of a spoken narrative (Bourguignon et al., 2020), presumably because auditory regions restore this temporal speech-related signature from the seen trajectory of the lip movements. Given that the representation of speech signals in temporally-aligned neural activity is essential for comprehension (Brodbeck and Simon, 2020; Giraud and Poeppel, 2012; Obleser and Kayser, 2019), this can be seen as indirect evidence that the auditory system supports lip reading by restoring key signatures of the underlying acoustic information based on the visual input.

Another view is that the visual system directly contributes to establishing speech representations separately from those established along the auditory pathway (Bernstein et al., 2011; O’Sullivan et al., 2017; Ozker et al., 2018). Visual speech contains temporal information that can be predictive of subsequent acoustic signals and allows mapping visual cues onto phonological representations (Campbell, 2008; Lazard and Giraud, 2017). Importantly, the visual cortex tracks dynamic lip signals (Park et al., 2016) and, as suggested recently, may also restore the unheard acoustic envelope of visually presented speech (Hauswald et al., 2018; Suess et al., 2022). Importantly, the evidence that visual speech induces information about the unheard speech acoustics along both auditory and the visual pathways may not be mutually exclusive, as both may contribute to a supramodal frame of reference for speech (Arnal et al., 2009; Rauschecker, 2012).

Many questions concerning a potential duality of acoustic speech-related representations during silent lip reading remain open (Bernstein and Liebenthal, 2014). First, each putative representation (in auditory and in visual cortex) was reported in a separate study and may have emerged mainly due to the specific stimuli or the specific analysis approach that was being applied (Bourguignon et al., 2020; Hauswald et al., 2018). Hence, it remains unclear whether both auditory and visual regions reflect acoustic speech features in parallel. Second, previous studies mainly capitalized on one-dimensional characterizations of the relevant sensory signals, such as the broadband speech envelope or the lip aperture. However, the cerebral encoding of speech is intrinsically multidimensional, and reflects temporal acoustic features such as the overall envelope or its derivative and spectral features such as pitch (Metzger et al., 2020; O’Sullivan et al., 2017; Oganian and Chang, 2019; Teoh et al., 2019). This raises the question of whether the previously observed restoration of acoustic speech-related information is tied to specific features, i.e. whether auditory regions preferentially encode the speech envelope or spectral features during lip reading. Third, it remains unclear whether a genuine acoustic feature is indeed represented independently of the physically observed lip movements and vice-versa. Alternatively, it may be previously reported restoration effects are largely explained by encoding of amodal information shared between both visual and acoustic modalities, which could be relayed to early sensory regions mainly by top-down processes. And finally, the behavioral relevance of the cerebral encoding of auditory speech features during lip reading remains unclear, as previous work mostly focused on neural signals but did not obtain direct measures of speech perception in natural language at the same time.

We systematically probed dynamic representations of acoustic and visual speech-related features in temporal and occipital brain regions during listening and viewing speech in the same participants using a mutual information approach (Daube et al., 2019; Keitel et al., 2018). This allowed us to provide a comprehensive assessment of how well temporal and occipital regions reflect either acoustic speech features or information about the lip trajectory, independently of each other, and both during hearing purely acoustic speech or while only seeing the speaker (lip reading). We probed the four main questions outlined above and found that both regions reflect unheard acoustic speech-related features independently of the physically observed lip movements. This ‘restoration’ of acoustic information in the temporal, but not the occipital, cortex was predictive of comprehension performance across participants.

## 2. Materials and Methods

The data analyzed in this study has been collected and analyzed in previous studies (Keitel et al., 2020, 2018). The analyses conducted here pose new questions and provide novel results beyond the previous work.

### 2.1 Participants and data acquisition

Data was collected from 20 native English speaking participants (9 female, age 23.6 ± 5.8 years mean ± SD). Due to prominent environmental artefacts in the MEG recordings, data from two participants were excluded from further analysis. Thus, the analyzed data is from 18 participants (7 female). All participants were screened for hearing impairment prior to data collection (Koike et al., 1994), had normal or corrected-to-normal vision and were all right-handed (Oldfield, 1971). All participants provided written informed consent and received monetary compensation of 10£/h. The experiment was approved by the College of Science and Engineering, University of Glasgow (approval number 300140078) and conducted in compliance with the Declaration of Helsinki.

MEG data was collected using a 248-magnetometer whole-head MEG system (MAGNES 3600 WH, 4-D Neuroimaging) with a sample rate of 1 kHz. Head positions were measured at the beginning and end of each run, using five coils placed on the participants’ heads. Coil positions were co-digitized with the participant’s head-shape (FASTRAK^®^, Polhemus Inc., VT, USA). Participants were seated in an upright position in front of a screen. Visual stimuli were displayed with a DLP projector at 25 frames per second, a resolution of 1280 × 720 pixels, and covered a visual field of 25 × 19 degrees. Acoustic stimuli were transmitted binaurally through plastic earpieces and 370-cm long plastic tubes connected to a sound pressure transducer and were presented in stereo at a sampling rate of 22,050 Hz.

### 2.2 Stimulus material

The stimulus material comprised two structurally equivalent sets of 90 unique matrix-style English sentences. Each sentence was constructed with the same sequence of linguistic elements, the order of which can be described with the following pattern [filler phrase, time phrase, name, verb, numeral, adjective, noun]. One such sentence for example was ‘I forgot to mention (filler phrase), last Thursday morning (time phrase) Marry (name) obtained (verb) four (numeral) beautiful (adjective) journals (noun)’. For each element, a list of 18 different options was created and sentences were constructed so that each single element was repeated ten times. Sentence elements were randomly combined within each set of 90 sentences. To measure comprehension performance for each sentence, a target word was defined in each sentence: either the adjective (first set of sentences) or the numeral (second set). Sentences lasted on average 5.4 ± 0.4 s (mean ± SD, ranging from 4.6 s to 6.5 s) and lasted a total of approximately 22 minutes. The speech material was spoken by a male British actor, who was tasked to speak clearly and naturally and to move as little as possible while speaking to assure that the lips center stayed at the same place in each video frame. Audiovisual recordings were gathered with a high-performance camcorder (Sony PMW-EX1) and an external microphone in a sound attenuating booth.

Participants were presented with audio-only (A), audiovisual (AV) or visual-only (V) speech material in three conditions (Keitel et al., 2018). However, for the present analysis we only focus on the A and V conditions, as in these one can best dissociate visual- and auditory-related speech representations given that only one physical stimulus was present. Furthermore, during the AV condition comprehension performance was near-ceiling (Keitel et al., 2020), making it difficult to link cerebral and behavioral data. To match the behavioral performance in the A and V condition, the acoustic speech was embedded in environmental noise. The noise for each trial was generated by randomly selecting 50 individual sounds from a set of sounds recorded from natural, everyday sources or scenes (e.g. car horns, talking people, traffic). For each participant the individual noise level was adjusted, as described previously (Keitel et al., 2020).

### 2.3 Experimental Design

Each participant was presented with each of the 180 sentences in three conditions (A, V and AV). The order of the conditions was fixed for all participants as A, AV and then V. Each condition was divided into 4 blocks of 45 sentences each, with two blocks being ‘adjective’ and two ‘number’ blocks. For each participant, the order of sentences within each block was randomized. The first sentence of each block was a ‘dummy’ trial that was subsequently excluded from analysis. During each trial, participants either fixated a dot (in A condition) or a small cross overlaid onto the mouth of the speaker’s face (in V condition). In the A condition, each sentence was presented as the respective audio recording, i.e. the spoken sentence, together with the background noise. In the V condition, only the video of the speaker’s face was presented and no sound was present. After each trial, four response option words (either adjectives or written numbers) were presented on the screen and participants had to indicate using a button press which word they had perceived. Inter-trial intervals were set to last about two seconds.

### 2.4 Preprocessing of stimulus material

From the stimulus material we extracted the following auditory and visual features. In the auditory domain, we derived the broadband envelope, the slope of the broadband envelope and the pitch contour. To derive the broadband envelope we filtered the acoustic waveform into twelve logarithmically spaced bands between 0.1 and 10 kHz (zero-phase 3rd order Butterworth filter with boundaries: 0.1, 0.22, 0.4, 0.68, 1.1, 1.7, 2.7, 4.2, 6.5, 10 kHz) and subsequently took the absolute value of the Hilbert transform for each band (Bröhl and Kayser, 2020). The broadband amplitude envelope was then derived by taking the average across all twelve band-limited envelopes and was subsequently down-sampled to 50 Hz. We computed the slope of this broadband envelope by taking its first derivative (Oganian and Chang, 2019). To characterize the pitch contour we extracted the fundamental frequency over time using the Praat software (‘to Pitch’ method with predefined parameters) (Boersma and van Heuven, 2001). This was done using the original acoustic waveform at a sampling rate of 22,050 Hz. The resulting pitch contour was again down sampled to 50 Hz. All three acoustic features together are labelled *AudFeat* in the following.

From the video recordings we derived the horizontal and vertical opening of the lips, the area covered by the lip opening, and its derivative. The lips were extracted based on the color of the lips in the video material using a custom-made algorithm. From these we determined the contour of the lip opening based on luminance values and deriving connected components from these (Giordano et al., 2017). The results were visually inspected to ensure accurate tracking of the lips. From this segmentation of the lip opening we derived the total opening (in pixels) and estimates of the respective diameters along the horizontal and vertical axes: these were defined between the outermost points along the respective horizontal (vertical) axis. These signals were initially sampled at the video rate of 30 fps. As for the auditory features, we computed the slope of the lip area. The time series of these visual features were then linearly interpolated to a sample rate of 50 Hz. Because the horizontal and vertical mouth openings are partially correlated with each other and with the total mouth opening, we selected the total area and the horizontal width as signals of interest, as the latter is specifically informative about the acoustic formant structure (Plass et al., 2020). We grouped the total lip area, it’s temporal derivative and the lip-width as signatures of lip features (*LipFeat*), which are of the same dimensionality as the acoustic features (AudFeat) described above.

For each of these features we derived its power spectrum and cross-coherence with the other features using MATLAB’s ‘pwelch’ and ‘mscoher’ functions using a window length of 3 s with 50% overlap and otherwise predefined parameters. The resulting spectra were log transformed and averaged across sentences. To visualize the cross-coherences we first obtained key frequency ranges of interest from our main results (c.f. Fig. 3) and averaged the coherences within two ranges of interest (0.5 - 1 Hz and 1 - 3 Hz). This was done to illustrate the stimuli’s spectral properties in the relevant frequency ranges.

### 2.5 MEG preprocessing

Preprocessing of MEG data was carried out using custom MATLAB scripts and the FieldTrip toolbox (Oostenveld et al., 2011). Each experimental block was processed separately. Individual trials were extracted from continuous data starting 2 s before sound onset and until 10 s after sound onset. The MEG data were denoised using a reference signal. Known faulty channels (N=7) were removed. Trials with SQUID jumps (3.5% of trials) were detected and removed using FieldTrip procedures with a cut-off z-value of 30. Data were band-pass filtered between 0.2 and 150 Hz using a zero-phase 4th order Butterworth filter and subsequently down sampled to 300 Hz before further artefact rejection. Data were visually inspected to find noisy channels (4.37 ± 3.38 on average across blocks and participants) and trials (0.66 ± 1.03 on average across blocks and participants). Noise cleaning was performed using independent component analysis with 30 principal components (2.5 components removed on average). Data were further down sampled to 50 Hz and bandpass filtered between 0.8 and 30 Hz using a zero-phase 3rd order Butterworth filter for subsequent analysis.

### 2.6 MEG source reconstruction

Source reconstruction was performed using Fieldtrip, SPM8, and the Freesurfer toolbox based on T1-weighted structural magnetic resonance images (MRIs) for each participant. These were co-registered to the MEG coordinate system using a semi-automatic procedure (Gross et al., 2013; Keitel et al., 2017). MRIs were then segmented and linearly normalized to a template brain (MNI space). We projected sensor-level time series into source space using a frequency-specific linear constraint minimum variance (LCMV) beamformer (Van Veen et al., 1997) with a regularization parameter of 7% and optimal dipole orientation (singular value decomposition method). The grid points had a spacing of 6 mm, thus resulting in 12,337 points. For whole-brain analyses, a subset of grid points corresponding to cortical gray matter regions only was selected (using the AAL atlas, Tzourio-Mazoyer et al., 2002), yielding 6,490 points in total. Within these we defined auditory and visual regions of interest (ROI) based on the brainnetome atlas (Yu et al., 2011). The individual ROIs were chosen based on previous studies that demonstrate the encoding of acoustic and visual speech features in occipital and superior temporal regions (Di Liberto et al., 2018; Giordano et al., 2017; Keitel et al., 2020; Teng et al., 2018). As auditory ROI we included Brodmann area 41/42, caudal area 22 (A22c), rostral area 22 (A22r) and TE1.0 and TE1.2. As visual ROI we defined the middle occipital gyrus (mOccG), occipital polar gyrus (OPC), inferior occipital gyrus (iOccG) and the medial superior occipital gyrus (msOccG).

### 2.7 MEG analysis

Source reconstructed MEG data were analyzed using a mutual information (MI) framework (Ince et al., 2017). The analysis relies on the notion that a significant temporal relation between cerebral signal and sensory features is indicating the cerebral encoding (or tracking) of the respective features by temporally entrained brain activity (Bröhl and Kayser, 2020; Keitel et al., 2018; Park et al., 2016). In the following we use the term ‘tracking’ when referring to such putative cerebral representations characterized using MI (Obleser and Kayser, 2019). To quantify the tracking of a given stimulus feature, or of a feature group, we concatenated the trial-wise MEG data and features along the time dimension and filtered these (using 3rd order Butterworth IIR filters) into typical frequency bands used to study dynamic speech encoding: 0.5 - 1 Hz, 1 - 3 Hz, 2 - 4 Hz, 3 - 6 Hz and 4 - 8 Hz (and 0.5 - 8 Hz). These were chosen based on previous work (Bröhl and Kayser, 2020; Etard and Reichenbach, 2019; van Bree et al., 2020; Zuk et al., 2021). The first 500 ms of each sentence were discarded to remove the influence of the transient sound-onset response. To compute the MI between filtered MEG and stimulus features, we relied on a complex-valued representation of each signal, which allowed us to include both the amplitude and phase information in the analysis: we first derived the analytic signal of both the MEG and stimulus feature(s) using the Hilbert transform and then calculated the MI using the Gaussian copula approach including the real and imaginary part of the Hilbert signals (Daube et al., 2019; Ince et al., 2017).

In a first step, we used this framework to visualize the tracking of AudFeat and LipFeat within the entire source space. This was mainly done to assert that the predefined ROIs used for the subsequent analysis indeed covered the relevant tracking of these features (Fig. 2). This analysis relied on a frequency range from 0.5 to 8 Hz and a range of stimulus-to-brain lags from 60 to 140 ms after stimulus onset. For the main analysis, we quantified the tracking of auditory or visual features and their dependencies specifically in each ROI (Fig. 3,4,5). To facilitate these analyses, we first determined the optimal lags for each feature, ROI and frequency band, given that the encoding latencies may differ between features and regions (Giordano et al., 2017). For this we determined at the group-level and for each feature group (i.e. AudFeat and LipFeat) and for each ROI and frequency band the respective lag yielding the largest group-level MI value (across participants and both A only and V only trials): for this we probed a range of lags between 0 and 500 ms in 20 ms steps. For the subsequent analyses, we used these optimal lags and averaged MI values obtained in a time window of −60 to 60 ms around these lags (computed in 20 ms steps).

**Fig. 1.**
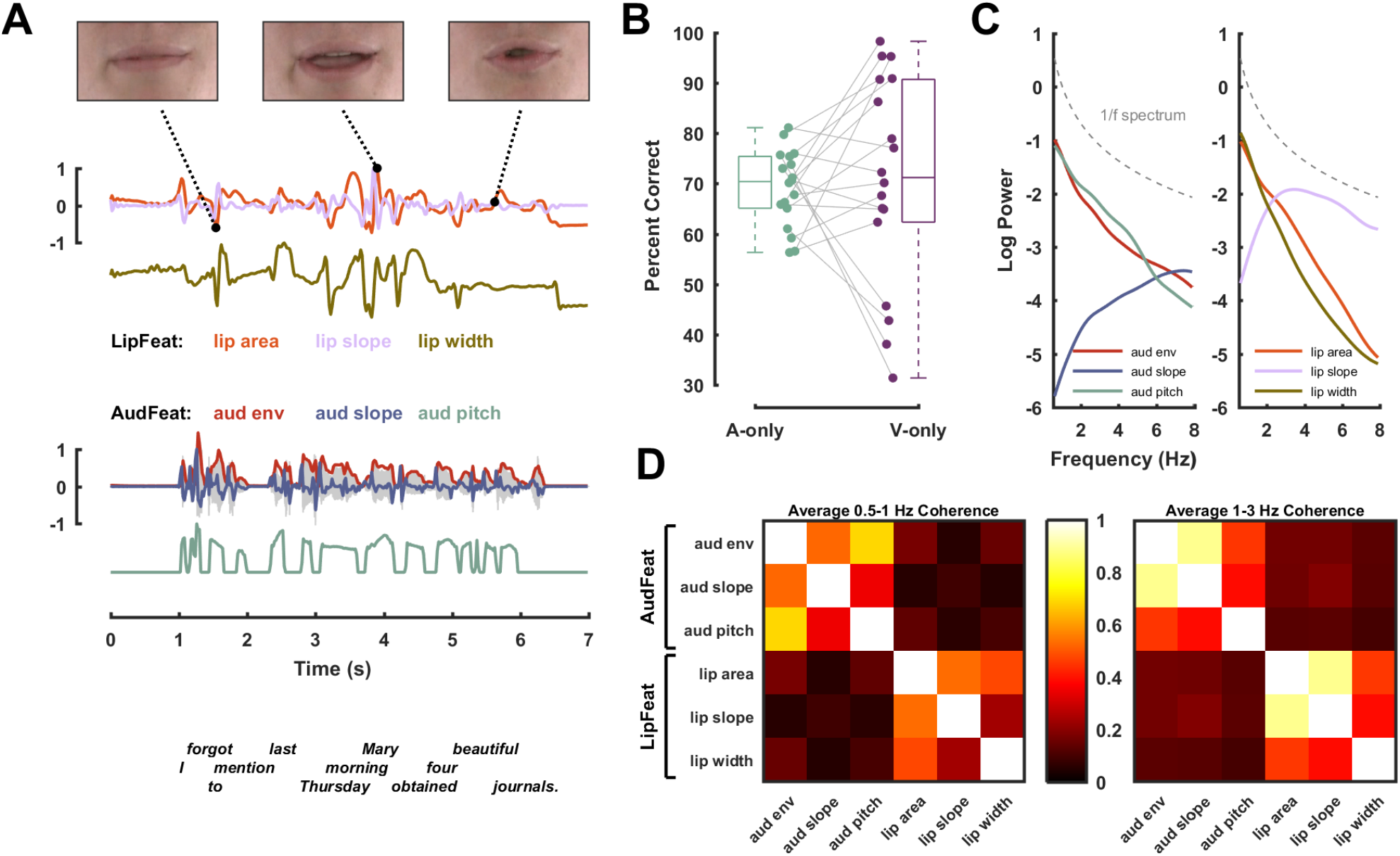
Stimulus material and experimental methodology. Acoustic and visual features were extracted from audiovisual speech material and were used to quantify their cerebral tracking during audio-only and visual-only presentations. (**A**) The stimulus material consisted of 180 audiovisual recordings of a trained actor speaking individual matrix-style English sentences. From video recordings we extracted three features describing the dynamics of the lip aperture: the area of lip opening (lip area), its slope (lip slope), and the width of lip opening (lip width); collectively termed ‘LipFeat’. The top row depicts three video frames illustrating the lip contour. From the audio waveform we extracted three acoustic features: the broadband envelope (aud env), its slope (aud slope), and a measure of dominant pitch (aud pitch); collectively termed ‘AudFeat’. (**B**) Trial-averaged percent correctly (PC) reported target words in auditory (A-only) and visual-only (V-only) conditions, with dots representing individual participants. (**C**) Logarithmic power spectra for individual stimulus features. For reference, a 1/f spectrum is shown as a dashed grey line. (**D**) Coherence between pairs of features averaged within two predefined frequency bands (0.5 - 1 Hz left; 1 - 3 Hz right, see Methods for details).

**Fig. 2.**
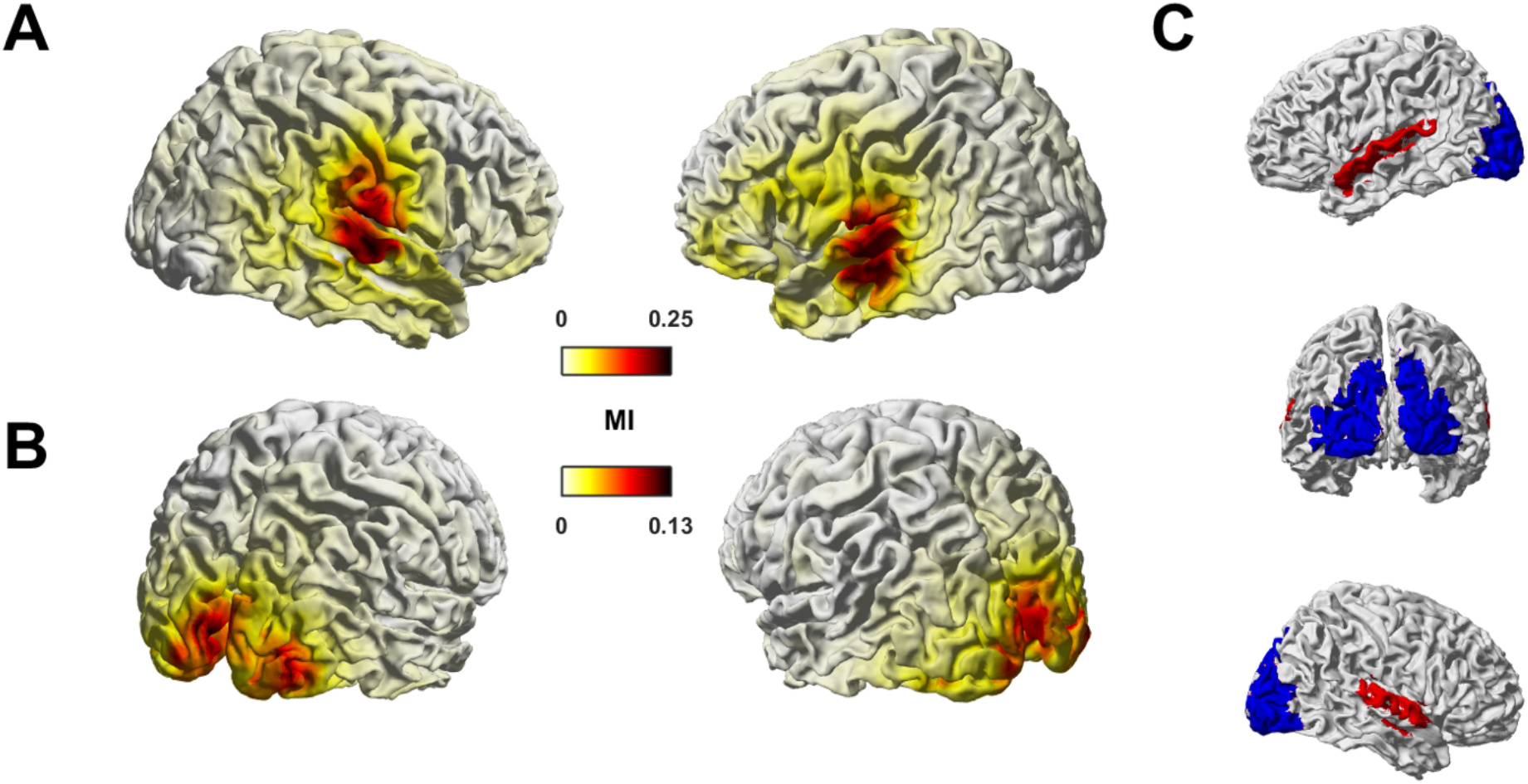
Tracking of auditory and visual features in MEG source space. The figure shows group-level median MI values for auditory (AudFeat; panel **A**) and lip features (LipFeat; panel **B**) in the frequency range from 0.5 - 8 Hz (n = 18 participants). (**C**) Colored shading indicates regions of interest: temporal regions in red include Brodmann area 41/42, caudal area 22 (A22c), rostral area 22 (A22r) and TE1.0 and TE1.2; occipital regions in blue include middle occipital gyrus (mOccG), occipital polar gyrus (OPC), inferior occipital gyrus (iOccG) and medial superior occipital gyrus (msOccG).

**Fig. 3.**
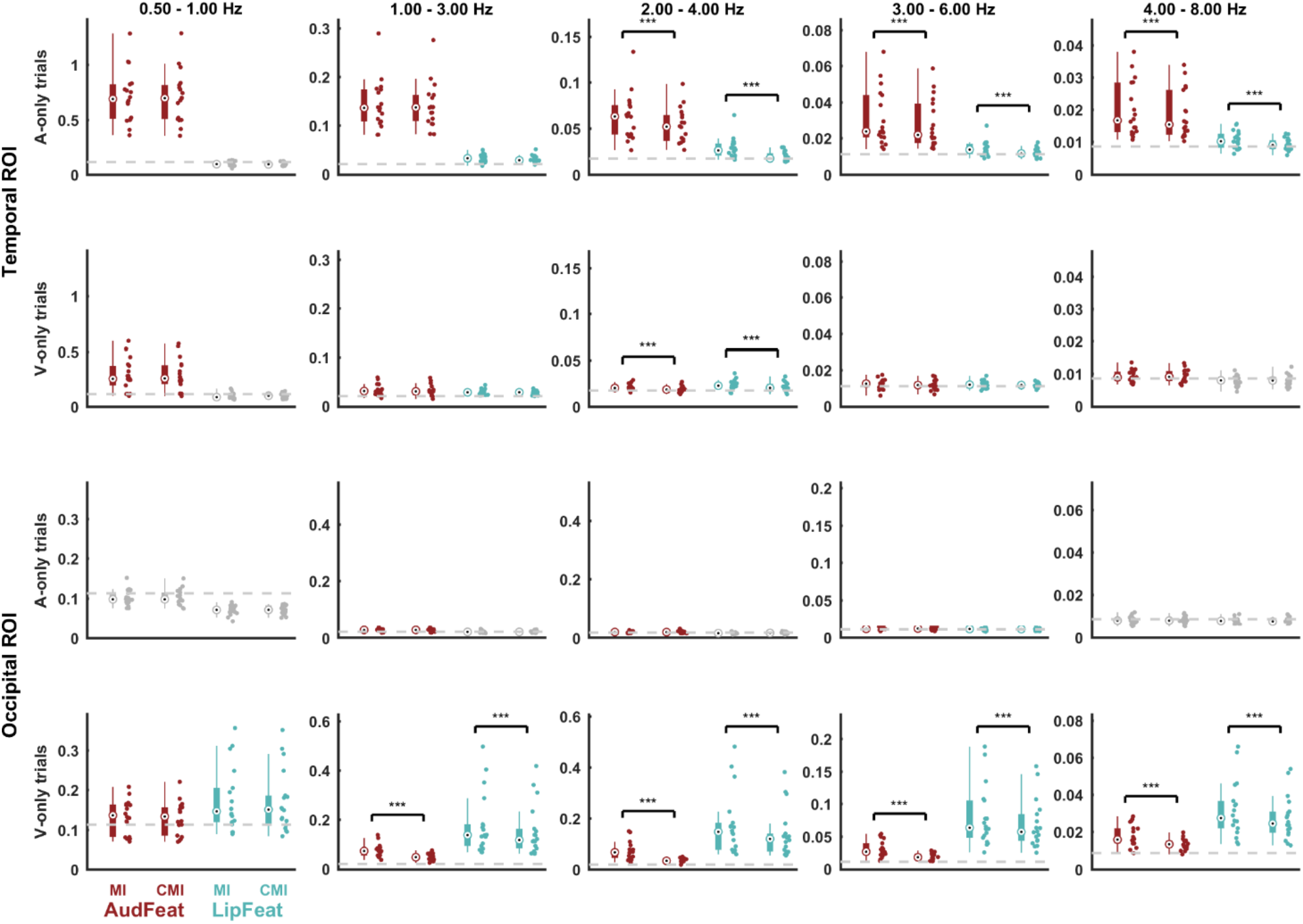
Feature tracking across regions of interest and conditions. For both conditions (A-only and V-only) and ROIs (temporal and occipital) the figure illustrates the strength of feature tracking for presented and physically not-present features (MI values) and the respective strength of tracking after partialling out the respective other feature group (CMI values). Each panel depicts (from left to right) the MI for AudFeat, the CMI for AudFeat partialling out LipFeat, the MI for LipFeat, and the CMI for LipFeat partialling out AudFeat. Dots represent individual participants (n = 18). Bars indicate the median, 25th and 75th percentile. The grey dashed line indicates the 99th percentile of the frequency-specific randomized maximum distribution correcting for all other dimensions. Conditions below a group-level significance threshold of 0.01 are greyed out. Brackets with asterisks indicate significant differences between MI and CMI values, based on a Wilcoxon signed-rank test (* p < 0.01, ** p < 0.005, *** p < 0.001). Units for MI and CMI are in bits.

**Fig. 4.**
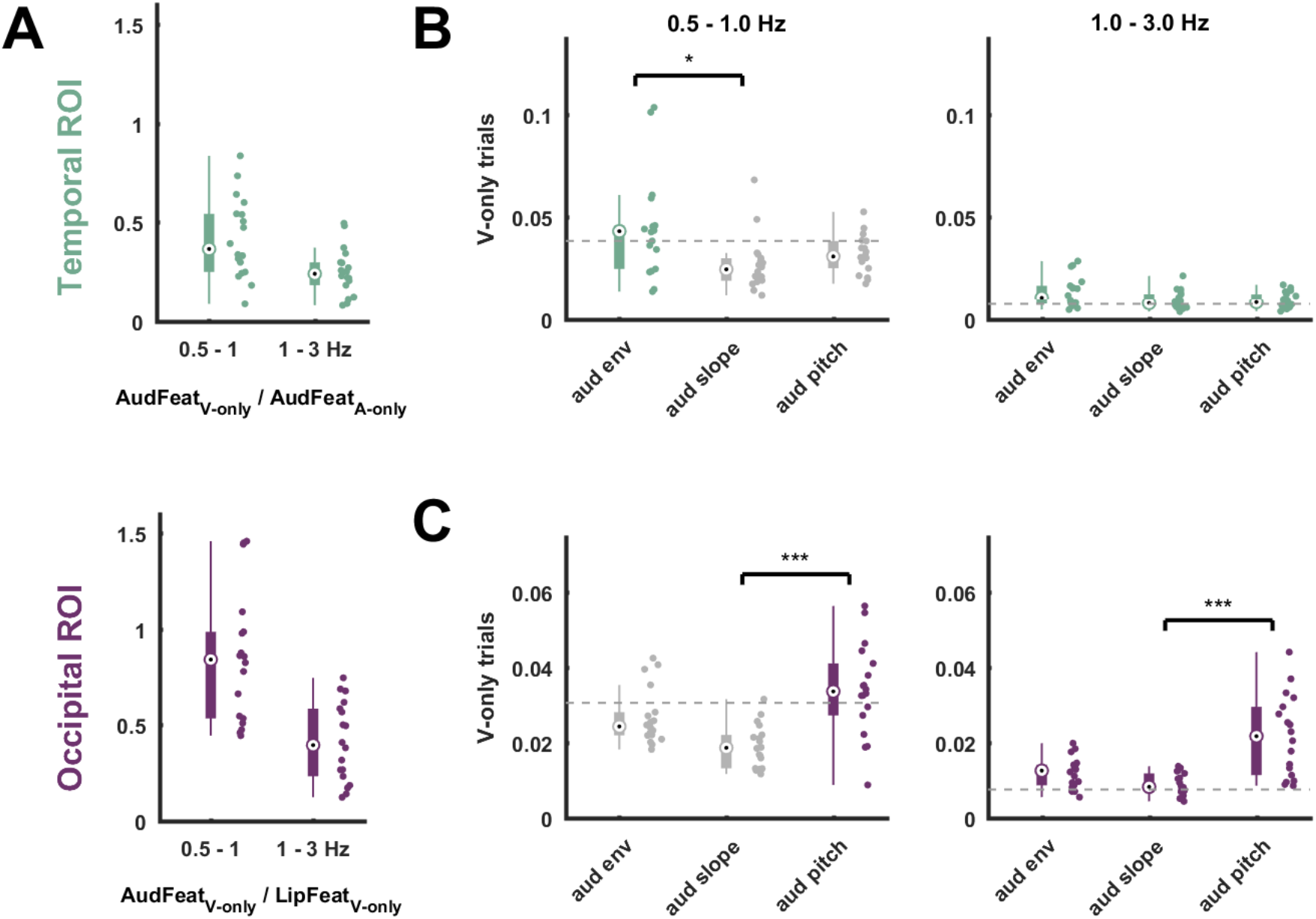
Modality dominance and tracking of individual auditory features during lip reading. (**A**) Comparison of the tracking of unheard AudFeat over the tracking of the modality-preferred sensory input in each ROI (i.e. AudFeat during A-only trials in the temporal ROI; LipFeat during V-only trials in the occipital ROI). (**B**,**C**) Tracking of individual auditory features during V-only trials conditioned on all other auditory and lip features in temporal (**B**) and occipital (**C**) ROIs. Brackets with asterisks indicate levels of significance from one-way Kruskal-Wallis rank test with post-hoc Tukey-Kramer testing (* p < 0.01, ** p < 0.005, *** p < 0.001). Dots represent individual data points. Bars indicate the median, 25th and 75th percentile. The grey dashed line indicates the 99th percentile of the frequency-specific randomized maximum distribution correction for all other features. Units in (**A**) are a ratio, in panels (**B**) and (**C**) units are in bits.

**Fig. 5.**
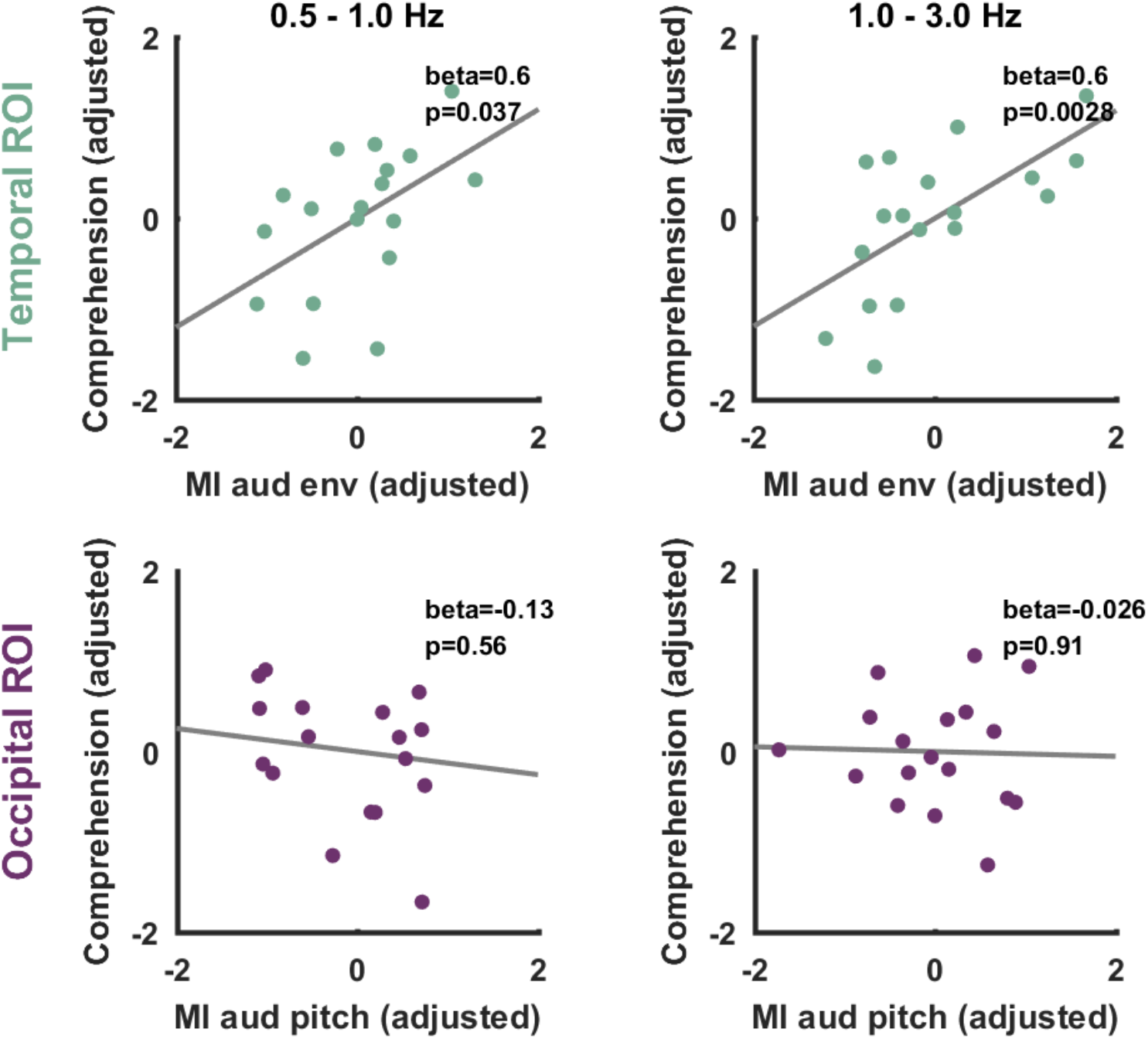
Association between lip reading performance and tracking of auditory features. Across participants the tracking of aud env during V-only trials in the temporal ROI but not the tracking of aud pitch in the occipital ROI was significantly associated with comprehension performance (PC) across participants in visual trials. Graphs show partial residual plots, dots represent individual data points and the line indicates the linear fit to the target variable from the full regression model.

Our hypotheses concerned both the MI between each feature group and the MEG and the statistical dependence of the tracking of each group on the tracking of the respective other group. To address this dependency, we determined whether the tracking of each feature group (in a given ROI and frequency band) is statistically redundant with (or possibly complementary to) the other group. For this we calculated the conditional MI between MEG and one feature group, partialling out the respective other group (Fig. 3, CMI values) (Giordano et al., 2017; Ince et al., 2017). Similarly, we also determined the conditional MI between the MEG and each individual feature, obtained by partialling out all other visual and auditory features (Fig. 4). To be able to compare the MI and CMI estimates directly, we ensured that both estimates had comparable statistical biases. To achieve this, we effectively derived the MI as a conditional estimate, in which we partialled out a statistically-unrelated variable. That is, we defined

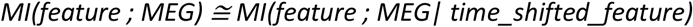

Here, time_shifted_feature is a representation of the respective feature(s) with a random time lag and hence no expected causal relation to the MEG. Each MI estimate was obtained by averaging this estimate over 2,000 repeats of a randomly generated time-shifted feature vector. To render the (conditional) MI estimates meaningful relative to the expectation of zero MI between MEG and stimulus features, we furthermore subtracted an estimate of the null-baseline of no systematic relation between signals. This was obtained by computing (conditional) MI values after randomly time-shifting the stimulus feature(s) and averaging the resulting surrogate MI estimates over 100 randomizations.

### 2.8 Relating MI to comprehension performance

The behavioral performance for each participant and condition was obtained as the percent correctly (PC) reported target words (obtained in a 4-choice task). To relate the tracking of specific features to comprehension performance, we accounted for potential spurious correlations between these due to the respective signal-to-noise ratio in each participants’ dataset. This was implemented using multiple regression, in which we predicted the PC in the visual trials based on i) the individual MI for *aud env* in the temporal ROI and the MI for *aud pitch* in the and occipital ROI as the primary variables of interest, and ii) the tracking of LipFeat (MI) in the occipital ROI in visual trials and iii) the tracking of AudFeat in the temporal ROI in auditory trials. The last two serve as potentially confounding variables, as they provide a proxy to the overall SNR of the speech and lip tracking in the respective dataset. By focusing on *aud env* / *aud pitch* in the temporal / occipital ROIs respectively, we predicted task performance based on the individual features that were most associated with the tracking of AudFeat (c.f. Fig. 4B,C). To establish these regression models, we z-scored the MI values of interest (variables i - iii) and the PC across participants. For the confounding variables, we applied the z-scoring for each frequency band and subsequently averaged the z-scored values across bands. For each frequency band, we created a single model containing all target and confounding variables. From the respective models we obtained the significance of each predictor of interest. Furthermore, we compared the predictive power of this full model with that of a reduced model not featuring the predictors of interest (variable i). From the likelihoods of each model we derived the relative Bayes factor (BF) between these based on the respective BIC values obtained from each model. For visualization we used partial residual plots using the procedure described by Velleman and Welsch (Velleman and Welsch, 1981). This procedure was applied to each individual feature of interest (i.e. *aud env* and *aud pitch*).

### 2.9 Statistical analysis

Statistical testing was based on a non-parametric randomization approach incorporating corrections for multiple comparisons (Nichols and Holmes, 2003). To test whether the group-level median MI (or CMI) values were significantly higher than expected based on the null hypothesis of no systematic temporal relation between sensory features and MEG, we proceeded in a similar fashion as in previous work (Bröhl and Kayser, 2020; Giordano et al., 2017): we obtained a distribution of 2,000 MI values between randomly time-shifted MEG and the stimulus vectors, while keeping the temporal relation of individual features to each other constant. This distribution was obtained for each participant, frequency band, feature (AudFeat and LipFeat), ROI (temporal, occipital) and condition (A-only, V-only) separately. To correct for multiple comparisons, we generated a maximum distribution across all dimensions except frequency bands, given that the MI values decreased considerably across bands (c.f. Fig. 3). We then tested the group-level median against the 99th percentile of this maximum distribution as a significance threshold, which effectively implements a one-sided randomization test at p < 0.01 corrected for all dimensions except frequency bands. To test for differences between MI and CMI values for a given condition, band and ROI, we also used a permutation approach combined with a Wilcoxon signed-rank test: first, we established the respective true Wilcoxon z-statistic between MI and CMI values; then we created a distribution of surrogate z-statistics under the null hypothesis of no systematic group-level effect, obtained by randomly permuting the labels of MI and CMI values 5,000 times. From this we obtained the maximum across features, bands, ROIs and conditions to correct for multiple comparisons and used the 99th percentile of this randomization distribution to determine the significance of individual tests.

The CMI values for individual features in Figure 4 were compared using a one-way repeated measure Kruskal-Wallis rank test, followed by a post-hoc Tukey Kramer multiple comparison. We used the same procedure to test for differences between CMI values in the sub-areas composing each ROI (Table 1). To test CMI values between hemispheres, we used a Wilcoxon signed rank test (Table 2). The resulting p-values were corrected for false discovery rate using the Benjamini-Hochberg procedure within each set of comparisons (Benjamini and Hochberg, 1995). In all tests an alpha level of α < 0.01 was deemed significant. For all statistical tests we provide exact p-values, except for randomization tests where the approximate p-values were smaller than the inverse of the number of randomizations.

**Table 1.** Feature tracking in individual anatomical areas within temporal and occipital ROIs. The table lists CMI values and a statistical comparison between the individual atlas-defined areas (Kruskal-Wallis tests, reporting chi-squares and p-values). Bold numbers indicate statistically significant results. P-values are FDR-corrected within this Table.

|  |  | 0.5 - 1 Hz |  |  |  | 1 - 3 Hz |  |  |  |
| --- | --- | --- | --- | --- | --- | --- | --- | --- | --- |
| ROI | Anatomical area | AudCMI | Chisq; pval | LipCMI | Chisq; pval | AudCMI | Chisq; pval | LipCMI | Chisq; pval |
| A-only trials |  |  |  |  |  |  |  |  |  |
| AC | A41/42 | 0.97 | 27.02 ; 4.7e-05 | 0.096 | 2.47 ; 0.59 | 0.19 | 29.62 ; 2.7e-05 | 0.032 | 5.14 ; 0.32 |
|  | TE1.0/1.2 | 0.56 |  | 0.099 |  | 0.11 |  | 0.028 |  |
|  | A22c | 0.86 |  | 0.098 |  | 0.18 |  | 0.033 |  |
|  | A22r | 0.5 |  | 0.093 |  | 0.095 |  | 0.029 |  |
| VC | mOccG | 0.1 | 3.50 ; 0.47 | 0.073 | 0.66 ; 0.88 | 0.025 | 2.71 ; 0.58 | 0.02 | 1.97 ; 0.66 |
|  | OPC | 0.09 |  | 0.069 |  | 0.025 |  | 0.02 |  |
|  | iOccG | 0.11 |  | 0.068 |  | 0.027 |  | 0.021 |  |
|  | msOccG | 0.11 |  | 0.074 |  | 0.029 |  | 0.021 |  |
| V-only trials |  |  |  |  |  |  |  |  |  |
| AC | A41/42 | 0.33 | 4.59 ; 0.36 | 0.1 | 5.14 ; 0.32 | 0.034 | 5.24 ; 0.32 | 0.027 | 1.00 ; 0.85 |
|  | TE1.0/1.2 | 0.25 |  | 0.088 |  | 0.031 |  | 0.028 |  |
|  | A22c | 0.34 |  | 0.1 |  | 0.035 |  | 0.027 |  |
|  | A22r | 0.22 |  | 0.085 |  | 0.028 |  | 0.029 |  |
| **VC** | mOccG | 0.13 | 3.90 ; 0.44 | 0.17 | 12.30 ; 0.026 | 0.045 | 8.20 ; 0.13 | 0.15 | 14.57 ; 0.012 |
|  | OPC | 0.14 |  | 0.19 |  | 0.06 |  | 0.2 |  |
|  | iOccG | 0.14 |  | 0.2 |  | 0.048 |  | 0.17 |  |
|  | msOccG | 0.11 |  | 0.11 |  | 0.039 |  | 0.082 |  |

**Tab. 2.** Feature tracking in each hemisphere. The table lists CMI values and a statistical comparison between hemispheres (Kruskal-Wallis tests, reporting chi-squares and p-values). P-values are FDR-corrected within this Table.

|  |  | 0.5 - 1 Hz |  |  |  | 1 - 3 Hz |  |  |  |
| --- | --- | --- | --- | --- | --- | --- | --- | --- | --- |
| ROI | Hemisphere | AudCMI | Z; pval | LipCMI | Z; pval | AudCMI | Z; pval | LipCMI | Z; pval |
| A-only trials |  |  |  |  |  |  |  |  |  |
| AC | left | 0.8 | 1.20 ; 0.59 | 0.094 | -0.33 ; 0.74 | 0.13 | -0.81 ; 0.59 | 0.028 | -1.11 ; 0.59 |
|  | right | 0.64 |  | 0.099 |  | 0.15 |  | 0.031 |  |
| VC | left | 0.1 | -0.37 ; 0.74 | 0.07 | -0.33 ; 0.74 | 0.027 | 0.81 ; 0.59 | 0.021 | 0.63 ; 0.65 |
|  | right | 0.1 |  | 0.072 |  | 0.025 |  | 0.02 |  |
| V-only trials |  |  |  |  |  |  |  |  |  |
| AC | left | 0.31 | 0.89 ; 0.59 | 0.098 | 0.76 ; 0.59 | 0.035 | 0.85 ; 0.59 | 0.025 | -1.85 ; 0.26 |
|  | right | 0.26 |  | 0.091 |  | 0.03 |  | 0.03 |  |
| VC | left | 0.11 | -2.24 ; 0.2 | 0.14 | -2.98 ; 0.046 | 0.043 | -1.68 ; 0.3 | 0.13 | -2.07 ; 0.21 |
|  | right | 0.15 |  | 0.2 |  | 0.054 |  | 0.18 |  |

## 3. Results

### 3.1 Acoustic and visual features are tracked in temporal and occipital cortices

Participants were presented with either spoken speech (A-only trials) or a silent video of the speaking face (V-only trials) and were asked to report a target word for each sentence in a 4-choice comprehension task. Previous work has shown that in this dataset temporal and occipital brain regions reflect auditory and visual speech signals respectively (Keitel et al., 2020). We extend this observation to the entire group of acoustic (AudFeat) or lip features (LipFeat) using a mutual information (MI) approach (Fig. 2). The whole-brain maps demonstrate the expected prevalence of acoustic (visual) tracking in temporal (occipital) regions. Given that our main questions concerned the tracking of features specifically in occipital and temporal brain regions, we focused the subsequent work on atlas-based regions of interest (Fig. 2; red and blue shaded areas).

### 3.2 Temporal and occipital cortex represent acoustic speech features during silent lip reading

To address the main questions of whether temporal and occipital cortices represent auditory and visual speech features during lip reading, we performed a comprehensive analysis of the tracking of both features across a range of frequency bands during auditory (A-only) and visual (V-only) conditions (MI values; Figure 3). Importantly, to determine whether the tracking of each feature group is possibly redundant with the tracking of the respective other feature group, we derived conditional MI values for each feature group, obtained by partialling out the respective other group (CMI values). By comparing MI and CMI values we can test, for example, whether the temporal ROI tracks the unheard speech envelope during silent lip reading also when discounting for the actually presented lip trajectory. In the following we discuss the results per sensory modality and region of interest.

As expected, when listening to speech (A-only), the temporal ROIs significantly track auditory features (AudFeat) in all frequency bands tested (Fig. 3, top row, red MI data; non-parametric randomization test, all bands: p < 5 x 10^-5^). This tracking persists when discounting potential contributions of the not-seen visual features (red CMI data all individually significant: p < 5 x 10^-5^), though in some bands the CMI values were significantly lower than the unconditional MI (Wilcoxon signed rank test comparing MI vs. CMI, 2 - 4 Hz: Z = 3.59, 3 - 6 Hz: Z = 3.68, 4 - 8 Hz: Z = 3.42, all comparisons: p < 2 x 10^-5^). During the same auditory trials, lip features are only marginally reflected in the temporal ROI, as shown by low but significant MI and CMI values above 1Hz (Fig. 3, top row, blue MI and CMI data; all bands above 1 Hz: p < 5 x 10^-5^). This tracking of visual features was significantly reduced when partialling out the physically presented auditory features (2 - 4 Hz: Z = 3.59, 3 - 6 Hz: Z = 3.68, 4 - 8 Hz: Z = 3.42, all comparisons: p < 2 x 10^-5^).

During lip reading (V-only), the temporal ROI tracks the unheard auditory features, particularly below 1 Hz (Fig. 3, 2nd row, red MI data; all bands: p < 5 x 10^-5^). Except in the 2 - 4 Hz range, the temporal ROI tracks the unheard AudFeat to a similar degree as when discounting the actually presented visual signal (significant red CMI values, all bands: p < 5 x 10^-5^): there were no significant differences between MI and CMI values except one band (2 - 4 Hz: Z = 3.42, p < 1 x 10^-4^, see asterisks). The physically presented lip movements during these V-only trials were also tracked significantly in the temporal ROI (Fig. 3, 2nd row; cyan MI and CMI data, 1 - 6 Hz: p < 5 x 10^-5^) but the CMI values were only marginally above chance level, suggesting that genuine visual representations in temporal regions are weak.

As expected, during lip reading (V-only) the occipital ROI tracks lip features (LipFeat) across frequency bands (Fig. 3, bottom row, cyan MI values; all bands: p < 5 x 10^-5^). Again, this tracking persists after partialling out the non-presented acoustic features (cyan CMI values; all bands: p < 5 x 10^-5^), although the CMI values were significantly lower than the MI (all bands above 1 Hz: Z ≥ 3.72, p < 2 x 10^-5^). This indicates some redundancy between the tracking of the physically present lip trajectory and that of the unheard auditory features. Confirming this, occipital tracking of the physically presented lip signals emerges in parallel with that of the non-presented auditory features (Fig. 3, bottom panel, red MI data; all bands: p < 5 x 10^-5^). This occipital tracking of unheard auditory features was significantly reduced when partialling out the lip signal (MI vs. CMI data; all bands above 1 Hz: Z ≥ 3.72, p < 2 x 10^-5^) but remained statistically significant (red CMI data; all bands: p < 5 x 10^-5^).

Finally, when listening to speech (A-only), the occipital ROI shows significant but weak tracking of auditory (Fig. 3, 3rd row, red MI data; 1 - 6 Hz: p < 5 x 10^-5^) and visual features (cyan MI data; only 3 - 6 Hz: p < 5 x 10^-5^), suggesting that purely acoustic signals have a weak influence on occipital brain regions.

Collectively, these results show the expected representations of auditory features in temporal cortex during listening to speech and of lip features in occipital cortex during lip reading. In addition, they reveal that during lip reading, both temporal and occipital regions represent unheard auditory features and do so independently of co-existing representations of the physically presented lip movements. In the auditory cortex this ‘restoration’ of auditory signals prevails in the low delta band (0.5 - 1 Hz), in the visual cortex this emerges in multiple bands.

To obtain an estimate of the effect size of the restoration of the unheard AudFeat during lip reading we expressed these CMI values relative to those of the tracking of the respectively modality-preferred inputs of each ROI (Fig. 4A): for temporal regions the tracking of AudFeat during A-only trials and for occipital regions the tracking of LipFeat during V-only trials. In the temporal ROI, the restoration effect size, i.e. the tracking of AudFeat during lip reading, was about a third as strong as this feature’s tracking while directly listening to speech (Fig. 4A; top row; AudFeat_V-only_ /AudFeat_A-only_; 0.5 - 1 Hz: median = 0.37, 1 - 3 Hz: median = 0.24). In the occipital ROI, the tracking of AudFeat was about half as strong or stronger compared to the tracking of lip features when seeing the speaker (Fig. 4A; bottom row; AudFeat_V-only_ /LipFeat_V-only_; 0.5 - 1 Hz: median = 0.84, 1 - 3 Hz: median = 0.4). Albeit smaller than the tracking of the respective modality-preferred sensory inputs, the restoration of unheard auditory features still results in a prominent signature in temporally aligned brain activity in both cortices.

### 3.3 Feature tracking is bilateral and prevails across anatomical brain areas

Having established the tracking of auditory and lip features in both temporal and occipital ROIs, we probed whether this tracking is possibly lateralized in a statistical sense and whether it potentially differs among the anatomical areas grouped into temporal and occipital ROIs respectively. For this analysis we focused on the conditional tracking of each feature group. Comparing CMI values among anatomical areas (averaged across hemispheres) for each ROI (occipital, temporal), frequency band (0.5 - 1 and 1 - 3 Hz), condition and feature group revealed a significant effect of area for AudFeat tracking in the temporal ROI during A-only trials (Table 1; 0.5 - 1 Hz: χ^2^(3) = 27.02, p = 4.7 x 10^-6^, ε^2^ = 0.35; 1 - 3 Hz: χ^2^(3) = 29.62, p = 2.7 x 10^-6^, ε^2^ = 0.39; p-values FDR-corrected). Post hoc comparisons revealed that in both bands, tracking of AudFeat was higher in A41/42 and A22c compared to TE1.0/1.2 and A22r (Tukey-Kramer test, all tests p < 10^-5^). The effect of Area was close to but not significant for LipFeat tracking in the occipital ROI during V-only trials (0.5 - 1 Hz: χ^2^(3) = 12.3, p = 0.026, ε^2^ = 0.14; 1 - 3 Hz: χ^2^(3) = 14.57, p = 0.012, ε^2^ = 0.17). Importantly, these results suggest that while the tracking of auditory features was stronger in the early auditory regions during A-only trials, the restoration of unheard auditory features during lip reading emerges to a similar degree among the individual temporal and occipital areas.

We performed a similar analysis comparing the CMI values within temporal or occipital ROIs between hemispheres. This revealed no significant effects of hemispheres (Table 2), hence offering no evidence for a statistical lateralization of feature tracking in the present data.

### 3.4 Occipital cortex reflects pitch more than other acoustic features during lipreading

Having established that occipital and temporal regions track unheard auditory features, we then asked how individual features contribute to these representations. For this we focused on the key condition of interest: the tracking of AudFeat in the delta range in V-only trials (Fig. 4 B,C). We quantified the CMI for each individual feature, while discounting the evidence about all other left-out visual and auditory features, hence focusing on the unique tracking of each individual acoustic feature.

For the temporal ROI this revealed the prominent tracking of aud env (Fig. 4B). In the in the 0.5 - 1 Hz band only the CMI for aud env was above chance (p < 5 x 10^-5^) and there was a significant effect of feature (Kruskal-Wallis rank test χ^2^(2) = 9.27, p = 9.1 x 10^-4^, ε^2^ = 0.14). Post-hoc tests revealed that the CMI for aud env differed significantly from that of aud slope (Tukey-Kramer test, p = 6.2 x 10^-4^; the other comparisons were not significant; p = 0.35 for env vs. slope and p = 0.22 for slope vs. pitch). In the 1 - 3 Hz band, the tracking of all auditory features was significant (all features: p < 5 x 10^-5^) and there was no significant effect of features (χ^2^(2) = 4.14, p = 0.13, ε^2^ = 0.04).

For the occipital ROI, this revealed a dominance of aud pitch (Fig. 4C). In the 0.5 - 1 Hz band, only the CMI of aud pitch was above chance (p < 5 x 10^-5^), a direct comparison revealed a significant effect of features (0.5 - 1 Hz: χ^2^(2) = 18.28, p = 1.07 x 10^-4^, ε^2^ = 0.32) and post-hoc tests revealed a significant difference between aud pitch and aud slope (p = 7.03 x 10^-5^), while the other comparisons were not significant (p = 0.26 for pitch vs. env and p = 0.02 for env vs. slope). In the 1 - 3 Hz range, the tracking of all features was significant (all features: p < 5 x 10^-5^), there was a significant effect of features χ^2^(2) = 19.2, p = 6.77 x 10^-5^, ε^2^ = 0.34), and post-hoc tests revealed a significant difference between pitch and slope (p = 3.61 x 10^-5^), while the other comparisons were not significant (p = 0.05 for pitch vs. env and p = 0.12 for env vs. slope). Collectively these results suggest that the restoration of acoustic information in occipital regions emphasizes spectral features, while in temporal regions this emphasizes the temporal speech envelope.

### 3.5 Tracking of auditory features is associated with lip reading performance

Finally, we probed the relevance of the restoration of unheard auditory features during silent lip reading for comprehension. For this we probed the predictive power of the MI about specific auditory features in either ROI for comprehension performance during V-only trials (Fig. 5). We specifically focused on the tracking of aud env in the temporal ROI and of aud pitch in the occipital ROI as the dominant feature-specific representations (c.f. Fig. 4B,C). Using linear models we predicted comprehension scores across participants based on the tracking indices of interest and while discounting for potential confounds from differences in signal-to-noise ratio.

The results show that variations in comprehension scores are well predicted by the collective measures of feature tracking (0.5 - 1 Hz: R^2^ = 0.74, 1 - 3 Hz: R^2^ = 0.8). Importantly, the tracking of aud env in the temporal ROI was significantly predictive of lip reading performance (0.5 - 1 Hz: β = 0.6, p = 0.037; 1 - 3 Hz: aud env β = 0.6, p = 2.8 x 10^-4^), while tracking of pitch in the occipital ROI was not (0.5 - 1 Hz: β = −0.13, p = 0.56; 1 - 3 Hz: β = −0.026, p = 0.91). This conclusion is also supported by Bayes factors for the added predictive power of aud env and aud pitch to these models (aud env in the temporal ROI; 0.5 - 1 Hz: BF = 3.12; 1 - 3 Hz: BF = 26.34; aud pitch in the occipital ROI; 0.5 - 1 Hz BF = 0.3; 1 - 3 Hz BF = 0.24).

## 4. Discussion

Natural face-to-face speech is intrinsically multidimensional and provides the auditory and visual pathways with partly distinct acoustic and visual information. These pathways could in principle focus mainly on the processing of their modality-specific signals, effectively keeping the two input modalities largely separated. Yet, many studies highlight the intricate multisensory nature of speech-related representations in the brain, including multisensory convergence at early stages of the hierarchy (Bernstein and Liebenthal, 2014; Crosse et al., 2015; Schroeder et al., 2008; Schroeder and Lakatos, 2009) as well as in classically amodal speech regions (Keitel et al., 2020; Mégevand et al., 2020; Scott, 2019). However, as the present results point out, the auditory and visual pathways are also capable of ‘restoring’ information about an absent modality-specific speech component. While seeing a silent speaker, both auditory and visual cortices track the temporal dynamics of the speech envelope and spectral features respectively, in a manner that is independent on the physically presented lip movements. Importantly, these ‘restored’ representations of acoustic speech features relate to participants’ comprehension, suggesting that they may form a central component of silent lip reading.

### 4.1 Auditory and visual cortex reflect acoustic speech features during lip reading

We systematically quantified the tracking of auditory and visual speech features during unisensory auditory and visual (lip reading) conditions in dynamically entrained brain activity. As expected, this analysis confirmed that early auditory and visual regions reflect acoustic and visual signals respectively at the time scales of delta (< 4 Hz) and theta (4 - 8 Hz) band activity, in line with a large body of previous work (Aiken and Picton, 2008; Bauer et al., 2020; Doelling et al., 2014; Giraud and Poeppel, 2012; Haegens and Zion Golumbic, 2018; Obleser and Kayser, 2019). In addition, we found that during lip reading both regions contained significant information about the unheard auditory features, also when discounting for the physically presented lip movements. This representation of acoustic features prevailed in the low delta band in auditory and the delta and theta bands in the visual cortex. Interestingly, this representation emphasized the temporal speech envelope in auditory cortex and spectral features (i.e. pitch) in the visual cortex. These results not only support that both regions are active during lip reading (Besle et al., 2008; Calvert et al., 1997; Calvert and Campbell, 2003; Ludman et al., 2000; Luo et al., 2010), but directly show that they contain temporally and feature-specific representations derived from lip movements that are also relevant for comprehension.

These results advance our understanding of how the brain exploits lip movements for speech-related processes in a number of ways. The restoration of auditory features during silent lip reading had been suggested in two previous studies, one showing the coherence of temporal brain activity with the non-presented speech envelope (Bourguignon et al., 2020) and another showing the coherence between occipital activity and the envelope (Hauswald et al., 2018; Suess et al., 2022). Yet, these studies differed in their precise experimental designs, their statistical procedures revealing the ‘restoration’ effect, and did not probe a direct link to comprehension performance. The present data demonstrate that such tracking of auditory speech-derived features indeed emerges in parallel and in the same participants, and, importantly, predicts comprehension. This suggests that perceptually relevant and possibly linked mechanisms may underlie the simultaneous processing of visual speech along visual and auditory pathways. In addition, our data show that this restoration emerges across a wider range of time scales as reported before (Bourguignon et al., 2020), and also when discounting for the physically present lip signals. The latter is particularly important, as the mere coherence of dynamic brain activity with the acoustic speech envelope may otherwise simply reflect those aspects of the physically-present visual speech that is directly redundant with the acoustic domain (Daube et al., 2019). Finally, our data suggest that this restoration is largely bilateral and emerges across a number of anatomically-identified areas, suggesting that it forms a generic property of the respective pathways.

Based on the same dataset as analyzed here, we recently showed that the identity of task-relevant words can be classified from the activity in multiple brain regions during lip reading and listening to speech (Keitel et al., 2020). While this previous study suggested that lip reading is facilitated by processes in early visual regions, the respective analysis focused on lexical identity and did not consider the individual features that may carry or contribute to such lexical information. The present results hence complement our previous work by demonstrating the alignment of temporal and occipital activity to the dynamics of the lip contour and specific acoustic features.

### 4.2 Lip reading activates a network of occipital and temporal regions

Previous work has shown that lip movements activate a network of temporal, parietal and frontal regions (Bourguignon et al., 2020; Calvert et al., 1997; Capek et al., 2008; O’Sullivan et al., 2017; Ozker et al., 2018; Paulesu et al., 2003; Pekkola et al., 2005) and that both occipital and motor regions can align their activity to the dynamics of lip movements (Park et al., 2018, 2016). The present data substantiate this, but also show that the representation of the physically visible lip trajectory along visual pathways is accompanied by the representation of spectral acoustic features, a type of selectivity not directly revealed previously (Suess et al., 2022). Spectral features are vital for a variety of listening tasks (Albouy et al., 2020; Bröhl and Kayser, 2020; Ding and Simon, 2013; Tivadar et al., 2020, 2018), and oro-facial movements provide concise information about the spectral domain. Importantly, as shown recently, seeing the speaker’s mouth allows discriminating formant frequencies and provides a comprehension benefit particularly when spectral speech features are degraded (Plass et al., 2020). This suggests a direct and comprehension-relevant link between the dynamics of the lip contour and spectral speech features (Campbell, 2008). Hence, a representation of acoustic features during silent lip reading may underlie the mapping of lip movements onto phonological units such as visemes, a form of language-specific representation emerging along visual pathways (Nidiffer et al., 2021; O’Sullivan et al., 2017).

Our results corroborate the notion that multisensory speech reception is not contingent only on high-level and modality-neutral representations. Rather, they suggest that cross-modal correspondences between auditory and visual speech exist along a number of dimensions, including basic temporal properties (Bizley et al., 2016; Chandrasekaran et al., 2009) as well as mid-level features, such as pitch or visual object features, whose representation is traditionally considered to be modality specific (Crosse et al., 2015; Plass et al., 2020; Schroeder et al., 2008; Zion Golumbic et al., 2013). Previous work has debated whether visual speech is mainly encoded along the auditory pathways or whether occipital regions contribute genuine speech-specific representations (O’Sullivan et al., 2017; Ozker et al., 2018). Our results speak in favor of occipital regions supporting speech reception by establishing multiple forms of speech-related information, including those aligned with the acoustic domain revealed here, and those establishing visemic categories based on complementary visual signals (Nidiffer et al., 2021; Suess et al., 2022). Which precise occipital areas and by which patterns of connectivity they contribute to comprehension remains to be investigated, but both kinds of representations may well emerge from distinct temporal-occipital networks (Bernstein and Liebenthal, 2014). While visemic information may be driven by object-related lateral occipital regions, the more auditory-aligned representations such as the restoration of spectral signatures may be directly driven by the connectivity between occipital areas and superior temporal regions, which play a key role for audio-visual speech integration (Arnal et al., 2009; Lazard and Giraud, 2017).

In the auditory cortex, the alignment of neural activity to the unheard speech envelope may reflect the predictive influence of visual signals on guiding the excitability of auditory pathways via low frequency oscillations (Schroeder et al., 2008). This alignment of auditory cortical activity to attended or expected sounds is well documented and has been proposed as a cornerstone of multisensory speech integration in general (Lakatos et al., 2008; Schroeder and Lakatos, 2009; Stefanics et al., 2010). One hypothesis is that this alignment may facilitate the segmentation or parsing of the speech stream (Ding et al., 2016; Giraud and Poeppel, 2012; Meyer et al., 2017). In this light the restoration of the speech envelope during lip reading suggests that such segmentation processes along the auditory pathways align to the presumed or expected acoustic counterpart underlying the received visual signal. This process would then act in parallel to visemic analysis in the visual pathway, and imply central functions of both auditory and visual pathways in lip reading.

## 5. Conclusion

Lip reading induces representations of the dynamic lip contour along visual pathways. Our results show that the brain derives representations of acoustic speech features from this sensory input as well, reflecting a form of restoration of acoustic speech-related features in auditory and visual cortices. In the auditory cortex these restored representations are predictive of lip reading performance, suggesting that they may form a central component of multisensory comprehension benefits.

## Credit author statement

Conceptualization: A.K., C.K., Project administration: A.K., C.K., Funding acquisition: C.K., Methodology: F.B., A.K., C.K., Software: F.B., A.K., Formal Analysis: F.B., Investigation: F.B., Data Curation: F.B., Supervision: C.K., Writing Original Draft: F.B., C.K., Writing Review & Editing: F.B., A.K., C.K.

## Data and code availability

Data and code used in this study are publicly available on the Data Server of the University of Bielefeld (https://gitlab.ub.uni-bielefeld.de/felix.broehl/fb02).

## Acknowledgement

This work was supported by the UK Biotechnology and Biological Sciences Research Council (BBSRC, BB/L027534/1) and the European Research Council (ERC-2014-CoG; grant No 646657)

